# A mathematical model-based approach to optimize loading schemes of isometric resistance training sessions

**DOI:** 10.1101/2020.04.16.044578

**Authors:** Johannes L. Herold, Andreas Sommer

## Abstract

Individualized resistance training (RT) is necessary to optimize training results. A model-based optimization of loading schemes could provide valuable impulses for practitioners and complement the predominant manual program design by customizing the loading schemes to the trainee and the training goals. We compile a literature overview of model-based approaches used to simulate or optimize the response to single RT sessions or to longterm RT plans in terms of strength, power, muscle mass, or local muscular endurance by varying the loading scheme. To the best of our knowledge, contributions employing a predictive model to algorithmically optimize loading schemes for different training goals are nonexistent in the literature. Thus, we propose to set up optimal control problems as follows. For the underlying dynamics, we use a phenomenological model of the time course of maximum voluntary isometric contraction force. Then, we provide mathematical formulations of key performance indicators for loading schemes identified in sport science and use those as objective functionals or constraints. We then solve those optimal control problems using previously obtained parameter estimates for the elbow flexors. We discuss our choice of training goals, analyze the structure of the computed solutions, and give evidence of their real-life feasibility. The proposed optimization methodology is independent from the underlying model and can be transferred to more elaborate physiological models once suitable ones become available.

## Introduction

### Resistance training and model-based approaches

Resistance training (RT) is a popular choice among athletes, rehabilitation patients, or the general public to improve physical performance. Benefits of RT include increased muscular strength and endurance, improved body composition, or enhanced functional capacity and quality of life [47]. To optimize results, individualized RT is necessary [20]. Therefore, training variables as exercise selection, frequency, volume, or intensity are adjusted to the trainee and the training goals. These adjustments are commonly performed by the trainee or a coach via trial-and-error [18].

To complement such a manual decision-making, many research areas like chemical or mechanical engineering have adopted methods from scientific computing, e.g., modeling, simulation, and optimization. For this reason, scientific computing is often considered to be the third pillar of methodology in science next to theory and experiment [33]. Nevertheless, sport science and exercise physiology are only slowly realizing the potential of model-based approaches [7]. In particular, applications covering loading schemes for resistance training are very limited. We refer to the literature overview in the next section to justify this claim.

A model-based optimization of loading schemes for RT could provide valuable impulses for practitioners and complement the predominant manual program design. By calibrating the model to the trainee, individual parameters are obtained. Then, optimized RT programs could be computed specifically for this trainee, exercise, and training goal based on a key performance indicator (KPI) accessible in the model. Furthermore, a comparison of effective loading schemes in practice and algorithmically optimized loading schemes could help to identify the driving stimuli for adaptations, e.g., the contributions of mechanical loading, metabolic stress, and muscle damage to hypertrophic adaptations [39] or the effect of different mechanical stimuli on strength and power adaptations [19]. Moreover, RT programs could be designed to induce the same level of metabolic disturbances. This would allow to increase the comparability between training approaches, e.g., between blood flow restriction training and conventional training.

### Purpose

In this work, we provide a literature overview of model-based approaches used to simulate or optimize the response to single RT sessions or to longterm RT plans in terms of strength, power, muscle mass, or local muscular endurance by varying the loading scheme. To the best of our knowledge, contributions employing a predictive model to algorithmically optimize loading schemes for different training goals are nonexistent in the literature. Thus, we propose to set up optimal control problems as follows. For the underlying dynamics, we use a phenomenological model of the time course of maximum voluntary isometric contraction (MVIC) force. Then, we provide mathematical formulations of key performance indicators for loading schemes identified in sport science and use those as objective functionals or constraints. Those KPIs are the force-time integral, the time-under-tension (TUT), the accumulated fatigue defined as loss of MVIC force, and variants thereof. We then solve those optimal control problems using previously obtained parameter estimates for the elbow flexors. Last, we discuss our results, point out limitations, and give an outlook on further research.

### Literature overview

In the following, we provide an overview of model-based approaches used to simulate or optimize an individual’s response to single RT sessions or to longterm RT plans in terms of strength, power, muscle mass, or local muscular endurance by varying the loading scheme. We do not include work that is restricted to the biomechanical analysis of RT exercises, the description of muscular fatigue during RT, or general models of the training-performance relationship without a specific application to RT. We begin with defining prerequisites which are necessary for a model to be used with our approach.

### Model prerequisites

To enable a real-life application for practitioners, the model used should fulfill several criteria. First, the inputs of the model, which correspond to the training plan of the trainee, have to be interpretable for practitioners. As such, using quantities which reduce the dimensionality of the training input [43] is not desirable. For example, using only volume load (defined as weight × repetitions × sets) [10] to describe the loading scheme of an RT session provides no information about the intensity distribution and is therefore unsuitable. Second, the parameters of the model should be identifiable through commonly available measurement procedures, e.g., force measurements, to avoid an overly laborious model calibration. Third, due to the high number of possible training inputs, the model should be suitable for high-dimensional optimization, i.e., for derivative-based optimization [24]. Fourth, the model should allow to incorporate real-life constraints into the optimization problem, e.g., days or weeks off [38]. Last, the model should be assessed for its predictive ability. We classify a model as predictive if it has been fit to a subset of the available data and the resulting parameter estimates can be used to predict the remaining data. We emphasize this, as the terminology is sometimes used differently and models are already classified as predictive if they fit the whole dataset – a property we call descriptive. However, overparameterization or other model deficiencies might diminish the model’s ability to predict unknown datasets. Benzekry et al. [9], for example, demonstrated this issue illustratively for tumor growth modeling. Furthermore, fit and prediction should be evaluated by suitable measures [41] and should not be judged based on the plots alone, as those are heavily depending on the chosen visualization.

### Existing models

Banister et al. [8] introduced a systems model based on the assumption that each training load induces a negative effect (fatigue) and a positive effect (fitness) on performance. As the original paper can not be found easily, we refer to Calvert et al. [17] for a description of the model. The ordinary differential equation (ODE) model has been adopted for various settings and several modifications have been proposed. The model is commonly known as Banister model or Fitness-Fatigue model and predominantly given in a time-discrete formulation. Busso et al. [14, 15] fitted variants of the Banister model to data from Olympic weightlifters. The authors used weighted weekly training volume as input and clean and jerk performance as output and correlated the model components to different hormones. However, the predictive ability of the model was not tested, i.e., the whole dataset was used for fitting the model. Model variants were furthermore used by Philippe et al. [35] to describe the response of rats to resistance training. In a subsequent work, the authors used exponential growth functions for this purpose [36]. In both works, model prediction was not tested.

Mader [31, 32] developed an ODE model of the active adaptation and regulation of protein synthesis on a cellular level. The model uses intensity of the functional activity as input and gives protein mass as an indicator of functional capacity as the most important output. The model is able to describe supercompensation as well as overtraining, which is demonstrated by simulating different scenarios. An extended version of the model has been proposed by Ullmer and Mader [46]. None of the variants were experimentally validated.

Gatti et al. [22] computed training plans for shoulder rehabilitation by determining the optimal number of sets per exercise for increasing maximum isometric strength given a time constraint. Two different objective functions were examined and compared to current practice. No statements about training intensity were made.

Gacesa et al. [21] used a nonlinear dynamic system to separately fit fatigue data and muscular growth data of the triceps brachii. The predictive ability of the model was not tested.

Arandjelovic [2] introduced a model of neuromuscular adaption to resistance training. In this model, the so-called capability profile of an athlete is modified depending on the execution of an exercise. The author subsequently used simulations to examine the influence of using fixed loads or accommodating loads on the training stimulus. Furthermore, the author proposed a framework to calibrate the model from video data [5, 7]. The model was found to successfully predict performance in the bench press and the squat. Resistance training can then be adjusted via trial-and-error by inspecting the simulated adaptations. Additionally, Arandjelovic used the model to examine training strategies to overcome the sticking point of an arm curl [3], to examine the influence of externally supplied momentum on the hypertrophy stimulus of a shoulder lateral raise [6], and to examine different loading mechanisms of a Smith machine [4]. Although these three studies are mainly of biomechanical nature, we mention them here, as they specifically aim at increasing force or muscle mass by a model-based examination of possible adaptations.

Wisdom et al. [48] proposed ODE models of muscle adaptation to chronic overstretch, overload, understretch, and underload and compared those models to experimental data. The predictive ability of the models was not tested. Zhou et al. [50] used similar dynamics to describe hypertrophy and atrophy of a muscle fiber given as cross-sectional area with muscle activation level as input. After fitting their model to experimental data, the authors simulated muscle atrophy during a spaceflight and how different exercises could serve as countermeasures.

Torres et al. [44] extended an energy balance model to account for the hypertrophic effects of resistance training and used the model for simulation studies. Moreover, the model was fit to data from elderly subjects following a resistance training routine. Resistance training input is described via a single scaling variable and has no direct interpretation in terms of volume, intensity, or frequency.

Herold et al. [24] constructed and validated a model of the time course of maximum voluntary isometric contraction force. Exemplarily, the model was used to algorithmically maximize the force-time integral (FTI) of an isometric RT session. We use this model as the foundation of our work, as it is – to the best of our knowledge – the only one to be tested for its predictive ability, suitable for derivative-based optimization, and directly interpretable for practitioners in terms of RT input. However, as the model provides a phenomenological description of muscular fatigue for different loading schemes, it does not directly link the RT input to a physiological adaptation of the trainee. Additionally, there still exist research gaps concerning the exact stimuli and mechanisms of muscular adaptation. To circumvent these issues, we provide mathematical formulations of KPIs for loading schemes identified in sport science and accessible in the model. Those KPIs are the force-time integral, the time-under-tension, the accumulated fatigue defined as loss of MVIC force, and variants thereof.

## Materials and methods

In this section, we describe the model and the optimization problems. For readers with a focus away from mathematical modeling, simulation, and optimization, we provide a short textual summary and then invite them to directly proceed to the results section if desired.

### Textual summary

Previous work [24] allows us to predict how MVIC force of a muscle group decreases and recovers under isometric loading (Equation (1)). Using mathematical methods of optimal control, this enables us to compute optimized isometric RT sessions (Equation (2)) with respect to different trainings goals. These training goals are constructed from the force-time integral, time-under-tension, or fatigue (Equations (3) to (6)).

### Model

For our numerical experiments, we use a phenomenological model of the time course of maximum voluntary isometric contraction force. We state the ordinary differential equation system and give a short explanation of the components. For a detailed description of the model, we refer to the original paper [24].

The model describes the current MVIC force capacity

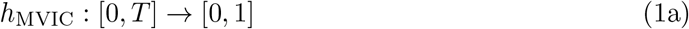

of a muscle (or muscle group) at joint level under an external isometric load

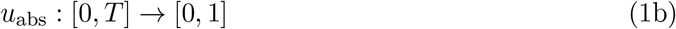

on the time horizon [0,*T*]. MVIC force capacity and external load are normalized to baseline MVIC force and are thus dimensionless. Moreover, the ranges of functions specified in this description are restricted to physiological meaningful values. The defining equations of the model are given as

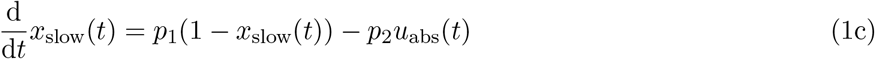

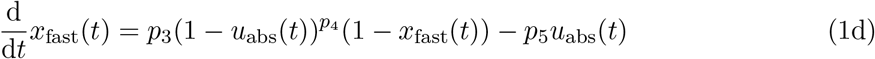

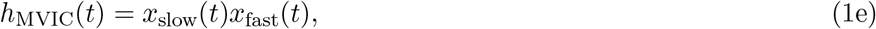

where

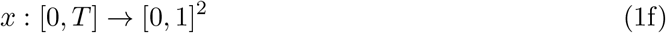

consists of two dimensionless state variables *x*_fast_ and *x*_slow_. The model furthermore contains five dimensionless parameters *p_i_* ∈ [0, ∞) for *i* ∈ {1,…,5} describing fatigue and recovery properties. The initial conditions for the states are given by

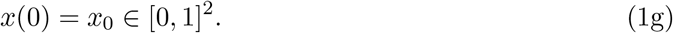

For an unfatigued muscle, one chooses *x*_0_ = (1,1)^⊤^. To simulate MVIC efforts, it is favorable to substitute

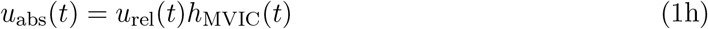

and use

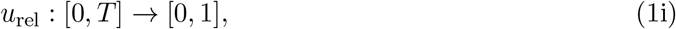

the load relative to the current force capacity, as input.

The model was validated with a comprehensive set of data from the elbow flexors [24]. We use the corresponding parameter estimates in this work.

### Optimal control problem

We use a multi-stage formulation on *n_s_* ≥ 2 stages – denoted by superscripts *i* ∈ {1,… *,n_s_*} – to model the resistance training sessions [24]. To include metrics for the TUT, the FTI, and the accumulated fatigue, we extend the model by three states tracking these quantities *x*_TUT_,*x*_FTI_, and *x*_fatigue_. The general multi-stage optimal control problem can then be formulated as

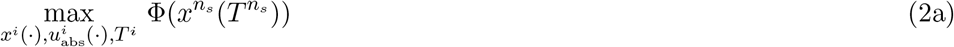

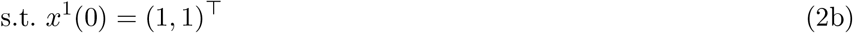

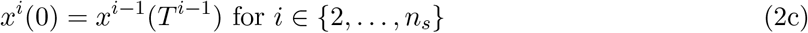

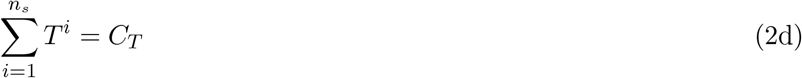

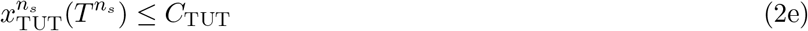

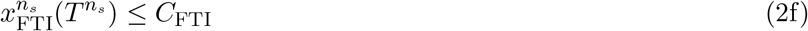

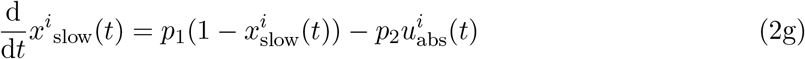

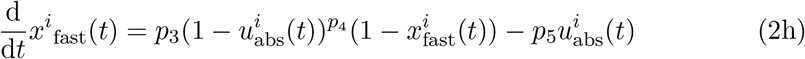

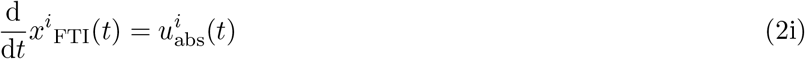

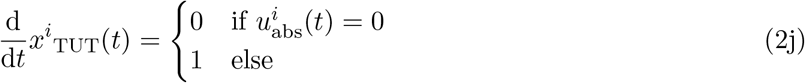

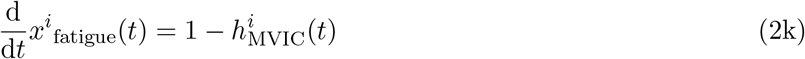

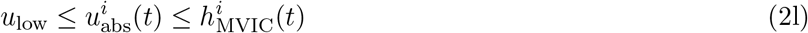

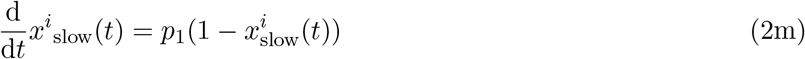

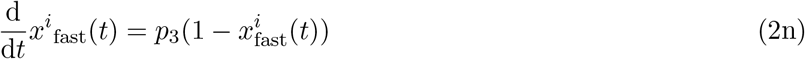

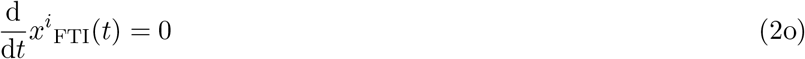

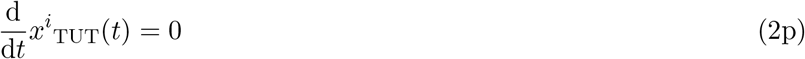

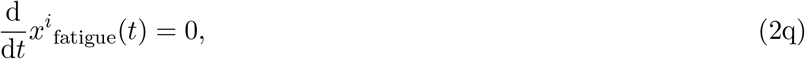

with *C_T_* being the total time and *C*_TUT_ and *C*_FTI_ the upper bounds on the total time-under-tension and the force-time integral. During odd numbered stages contractions with *u*_low_ ≤ *u*_abs_ are possible. Even numbered stages are considered rest periods. The duration *T*^i^ of each stage is being optimized. We adapt this optimal control problem to different scenarios in the following. If not mentioned otherwise, all sessions last 20 min, allow *n_c_* = 25 possible contractions and have no restrictions on FTI or TUT. This implies *C_T_* = 1200 s, *n_s_* = 49 and neglecting Constraints (2e) and (2f). Table 1 gives an overview of the symbols used in the problem formulation.

**Table 1.**
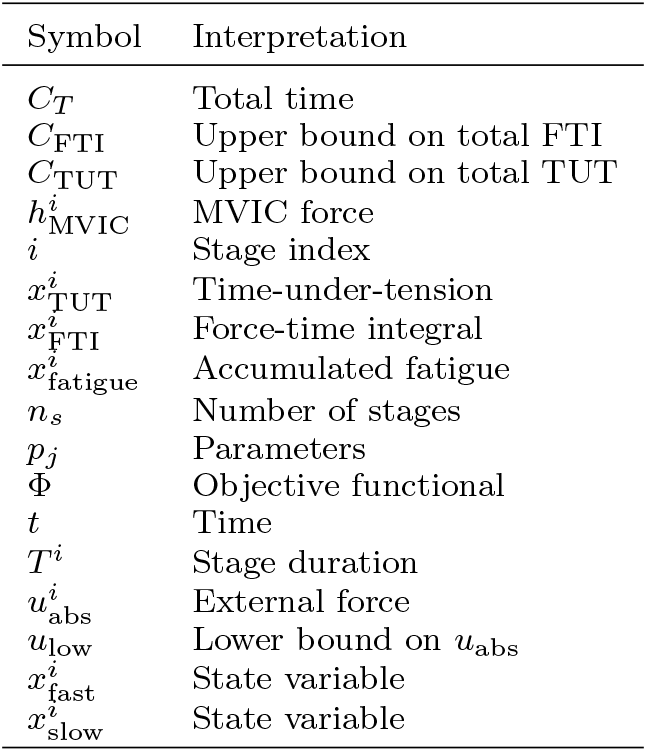
Overview of symbols used in the multi-stage optimal control problem (2).

To solve the problems numerically, we employ a first-discretize-then-optimize strategy. We use the optimal control software MUSCOD-II [28, 29], which originates from the work of Bock and Plitt [11] and implements a direct multiple shooting approach.

In the following, we present how this general optimal control problem formulation (2) is adapted to different sessions (labeled Session A to K). We refer to Table 2 for a concise overview.

**Table 2.**
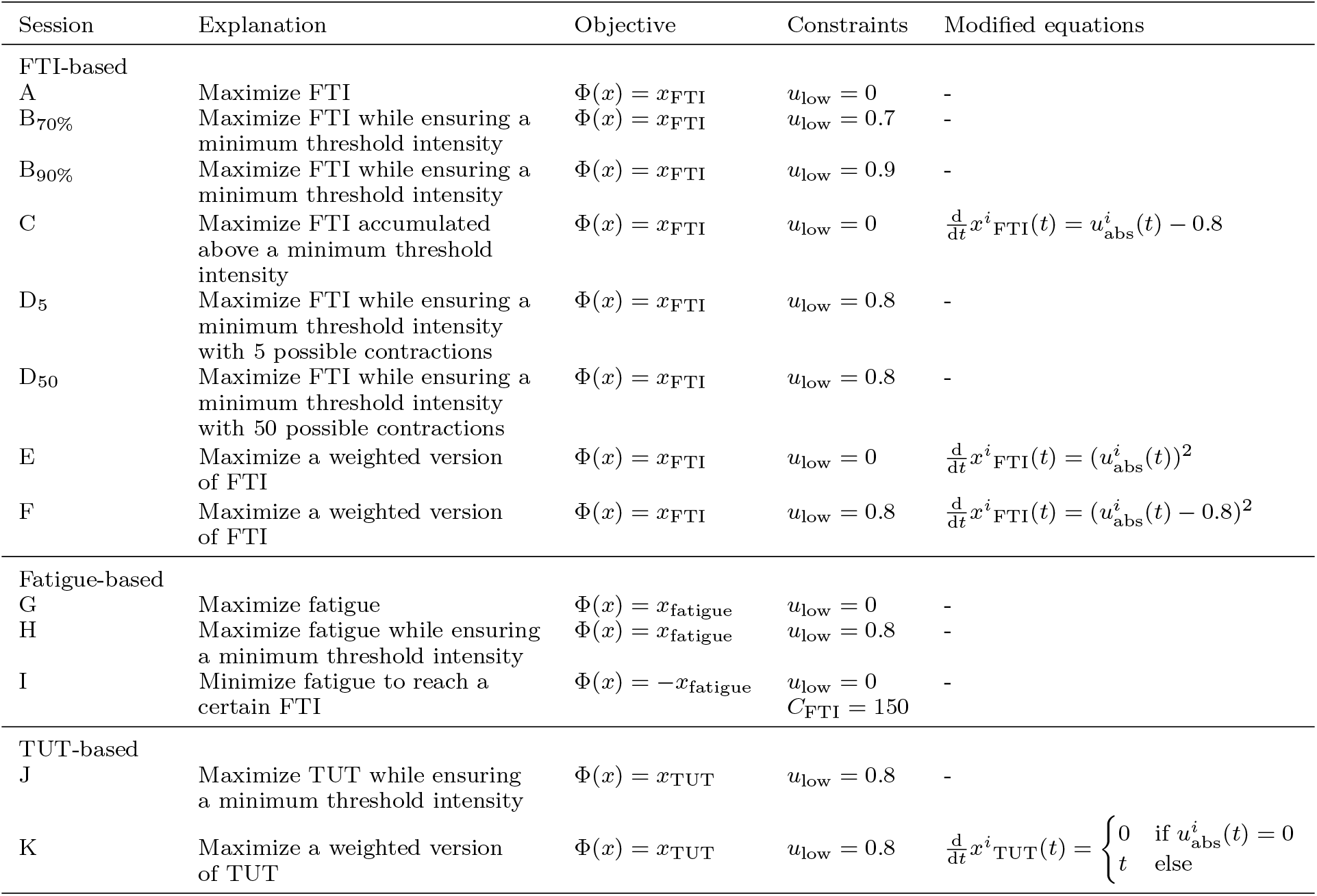
Overview of sessions used in this work. If not mentioned otherwise, all sessions last 20 min and allow 25 possible contractions.

| Session | Explanation | Objective | Constraints | Modified equations |
| --- | --- | --- | --- | --- |
| FTI-based |  |  |  |  |
| A | Maximize FTI | $\Phi(x) = x_{\text{FTI}}$ | $u_{\text{low}} = 0$ | - |
| B<sub>70%</sub> | Maximize FTI while ensuring a minimum threshold intensity | $\Phi(x) = x_{\text{FTI}}$ | $u_{\text{low}} = 0.7$ | - |
| B<sub>90%</sub> | Maximize FTI while ensuring a minimum threshold intensity | $\Phi(x) = x_{\text{FTI}}$ | $u_{\text{low}} = 0.9$ | - |
| C | Maximize FTI accumulated above a minimum threshold intensity | $\Phi(x) = x_{\text{FTI}}$ | $u_{\text{low}} = 0$ | $\frac{d}{dt}x^i_{\text{FTI}}(t) = u^i_{\text{abs}}(t) - 0.8$ |
| D<sub>5</sub> | Maximize FTI while ensuring a minimum threshold intensity with 5 possible contractions | $\Phi(x) = x_{\text{FTI}}$ | $u_{\text{low}} = 0.8$ | - |
| D<sub>50</sub> | Maximize FTI while ensuring a minimum threshold intensity with 50 possible contractions | $\Phi(x) = x_{\text{FTI}}$ | $u_{\text{low}} = 0.8$ | - |
| E | Maximize a weighted version of FTI | $\Phi(x) = x_{\text{FTI}}$ | $u_{\text{low}} = 0$ | $\frac{d}{dt}x^i_{\text{FTI}}(t) = (u^i_{\text{abs}}(t))^2$ |
| F | Maximize a weighted version of FTI | $\Phi(x) = x_{\text{FTI}}$ | $u_{\text{low}} = 0.8$ | $\frac{d}{dt}x^i_{\text{FTI}}(t) = (u^i_{\text{abs}}(t) - 0.8)^2$ |
| Fatigue-based |  |  |  |  |
| G | Maximize fatigue | $\Phi(x) = x_{\text{fatigue}}$ | $u_{\text{low}} = 0$ | - |
| H | Maximize fatigue while ensuring a minimum threshold intensity | $\Phi(x) = x_{\text{fatigue}}$ | $u_{\text{low}} = 0.8$ | - |
| I | Minimize fatigue to reach a certain FTI | $\Phi(x) = -x_{\text{fatigue}}$ | $u_{\text{low}} = 0$<br>$C_{\text{FTI}} = 150$ | - |
| TUT-based |  |  |  |  |
| J | Maximize TUT while ensuring a minimum threshold intensity | $\Phi(x) = x_{\text{TUT}}$ | $u_{\text{low}} = 0.8$ | - |
| K | Maximize a weighted version of TUT | $\Phi(x) = x_{\text{TUT}}$ | $u_{\text{low}} = 0.8$ | $\frac{d}{dt}x^i_{\text{TUT}}(t) = \begin{cases} 0 & \text{if } u^i_{\text{abs}}(t) = 0 \\ t & \text{else} \end{cases}$ |

### FTI-based goals

Resistance training volume is an important determinant of longterm adaptations [20]. For isometric contractions, where no actual physical work is performed, the force-time integral is an often used analogue of work [37]. Thus, for Session A, we maximize the FTI accumulated during an RT session without imposing restrictions on the contraction intensity, i.e., Φ(*x*) = *x*_FTI_ and *u*_low_ = 0.

To increase maximum strength, high loads are recommended by some researchers, e.g., by the American College of Sports Medicine [1]. Therefore, the model has previously been used to compute an exemplary optimized RT session, which maximizes the FTI and ensures that the contraction intensity is higher than a minimum threshold intensity of 80 % of baseline MVIC force [24]. We adopt this example and examine how lowering or raising the minimum threshold intensity influences the solution. For Session B_70%_, we set Φ(*x*) = *x*_FTI_ and *u*_low_ = 0.7. For Session B_90%_, we set Φ(*x*) = *x*_FTI_ and *u*_low_ = 0.9.

As an alternative to the full FTI maximized in Session A, one can use the FTI accumulated above the minimum threshold intensity as an indicator of effective training volume. For Session C, we thus set *u*_low_ = 0 and replace Equation (2i) with

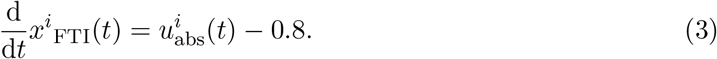

A similar measure has been used by Burnley [13] when examining work capacity above critical torque.

For Session D, we examine the influence of the number of possible contractions on Session B and compute the solution for *n_c_* ∈ {5,6,…, 49, 50} possible contractions. This allows to investigate if more but expectedly shorter contractions allow to accumulate a higher FTI while ensuring a minimum threshold intensity of *u*_low_ = 0.8 and if the additional possible contractions are actually realized in the solution.

Instead of choosing a minimum threshold intensity, we can emphasize higher loads by evaluating a weighting function on the integrand of the FTI. For demonstration purposes, we choose a quadratic weighting function for Session E. Therefore, we set Φ(*x*) = *x*_FTI_ and replace Equation (2i) with

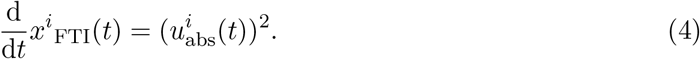

*u*_low_ is set to 0. A similar approach has been used by Arandjelovic [6] to describe the hypertrophy stimulus of a resistance training set, although he used a sigmoid function, which can be interpreted as a smoothing of the constraint *u*_low_ ≤ *u*_abs_ used in Session B.

A similar weighting can be applied to Session C by replacing Equation (2i) with

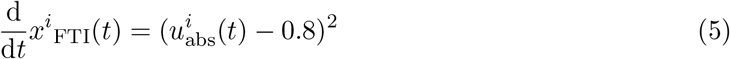

and setting the objective functional to Φ(*x*) = *x*_FTI_ for Session F. In contrast to Session C, *u*_low_ = 0.8 is necessary here, as otherwise *u*_abs_ = 0 would be the solution.

### Fatigue-based goals

Effects of fatigue, e.g., metabolic stress or increased motor unit recruitment, have been attributed to trigger and/or positively influence muscle hypertrophy [39]. We examine which loading scheme maximizes fatigue, defined as the accumulated loss of MVIC force over time. Thus, for Session G, we choose Φ(*x*) = *x*_fatigue_ and *u*_low_ = 0.

For Session H, we maximize fatigue while ensuring a minimum threshold intensity of 80 % of baseline MVIC force. Therefore, we choose Φ(*x*) = *x*_fatigue_ and *u*_low_ = 0.8.

In contrast to maximizing fatigue, it might also be desired to accumulate a certain amount of work while minimizing fatigue, e.g., during the tapering period before a competition. For Session I, we exemplarily choose Φ(*x*) = −*x*_fatigue_ and *C*_FTI_ = 150 s.

### TUT-based goals

Several authors have examined time-under-tension as a determinant of acute responses and longterm adaptations to RT (e.g., Burd et al. [12] or Schott et al. [40]). Therefore, for Session J, we maximize TUT while ensuring a minimum threshold intensity by choosing Φ(*x*) = *x*_TUT_ and *u*_low_ = 0.8.

Session J does not take into account the duration of the contractions used to accumulate the total TUT. However, some author have reported different adaptations to short and long duration contractions with greater hypertrophy occurring after long duration contractions [40]. Thus, to weight the duration of contractions quadratically, we replace Equation (2j) with

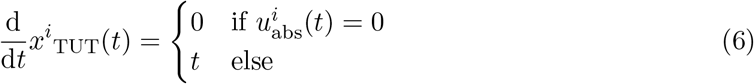

for Session K. All other settings are kept as in Session J.

## Results

In the following, we provide the results of our computations. Here, we focus on the structure of the computed solutions. For readers who skipped the methods section, we redescribe the scenarios without the mathematical details. We refer to Table 2 for a concise overview. If not mentioned otherwise, all sessions last 20 min and allow 25 possible contractions.

### FTI-based goals

Resistance training volume is an important determinant of longterm adaptations [20]. For isometric contractions, where no actual physical work is performed, the force-time integral is an often used analogue of work [37]. Thus, for Session A, we maximize the FTI accumulated during an RT session without imposing restrictions on the contraction intensity. Figure 1a illustrates the model response obtained by simulating Session A.

**Figure 1.**
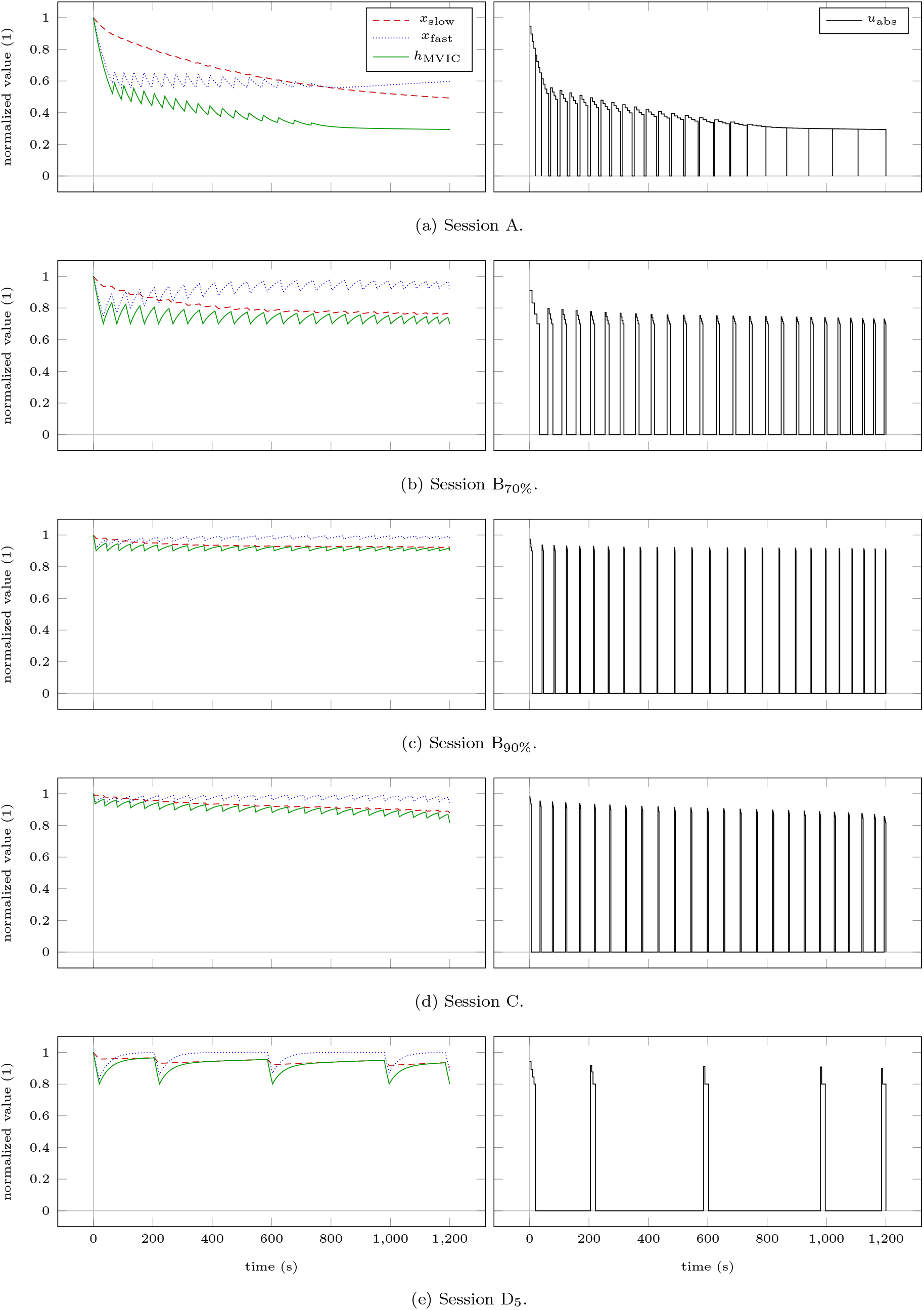
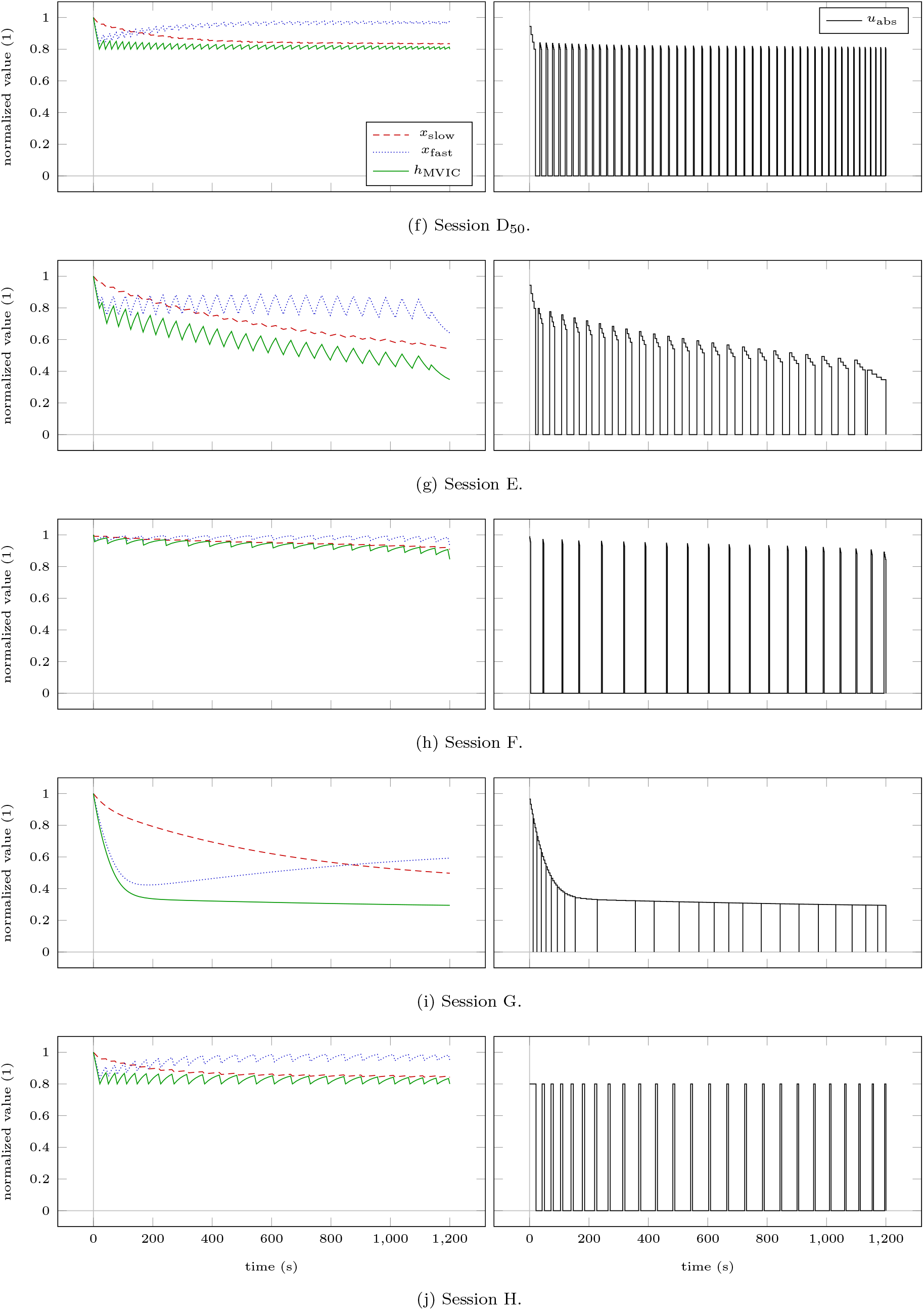
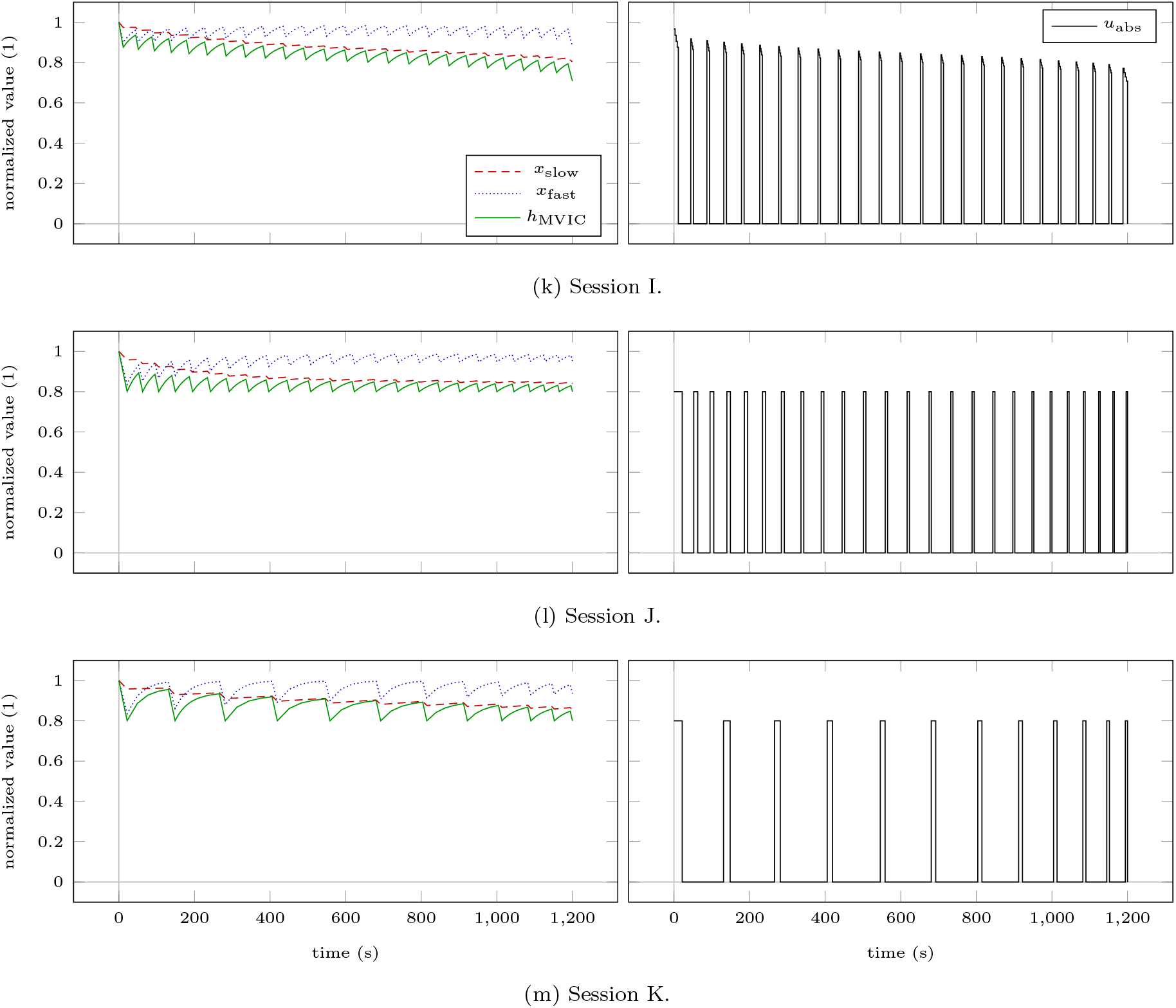
Model response obtained by simulating Sessions A to D_5_. We refer to the text and Table 2 for an explanation of the individual sessions. The left column depicts the model response. The absolute force input is illustrated in the right column. Model response obtained by simulating Sessions D_50_ to H. We refer to the text and Table 2 for an explanation of the individual sessions. The left column depicts the model response. The absolute force input is illustrated in the right column. Model response obtained by simulating Sessions I to K. We refer to the text and Table 2 for an explanation of the individual sessions. The left column depicts the model response. The absolute force input is illustrated in the right column.

To increase maximum strength, high loads are recommended by some researchers, e.g., by the American College of Sports Medicine [1]. Therefore, the model has previously been used to compute an exemplary optimized RT session, which maximizes the FTI and ensures that the contraction intensity is higher than a minimum threshold intensity of 80 % of baseline MVIC force [24]. We adopt this example and examine how lowering or raising the minimum threshold intensity to 70 % or 90 % of baseline MVIC force influences the structure of the solution. Figures 1b and 1c illustrate the model response obtained by simulating Sessions B_70%_ and B_90%_.

For Session C, as an alternative to the full FTI maximized in Session A, one can use the FTI accumulated above the minimum threshold intensity as an indicator of effective training volume. A similar measure has been used by Burnley [13] when examining work capacity above critical torque. Figure 1d illustrates the model response obtained by simulating Session C.

For Session D, we examine the influence of the number of possible contractions on Session B and compute the solution for 5 to 50 possible contractions. This allows to investigate if more but expectedly shorter contractions allow to accumulate a higher FTI while ensuring a minimum threshold intensity of 80 % of baseline MVIC force and if the additional possible contractions are actually realized in the solution. Figures 1e and 1f illustrate the model response obtained by simulating Sessions D_5_ and D_50_. Figure 2 depicts the objective functional value in dependency of the number of possible contractions. Figure 3 depicts the durations of contractions and rests in dependency of the number of possible contractions. For all sessions, all 25 possible contractions are realized.

**Figure 2.**
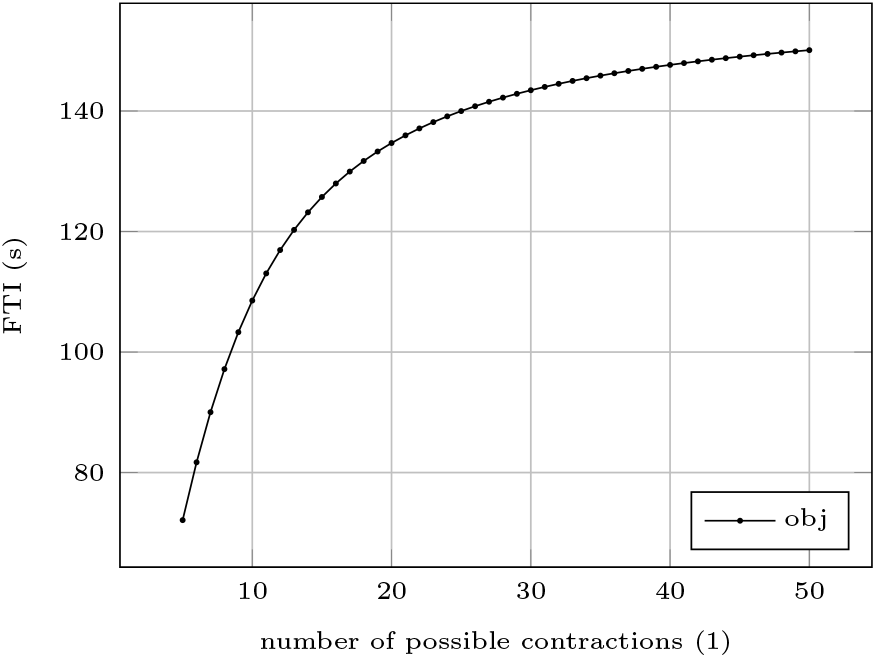
Dependency of the objective functional value on the number of possible contractions for Sessions D_5_ to D_50_. Increasing the number of possible contractions increases the FTI of the computed solution.

**Figure 3.**
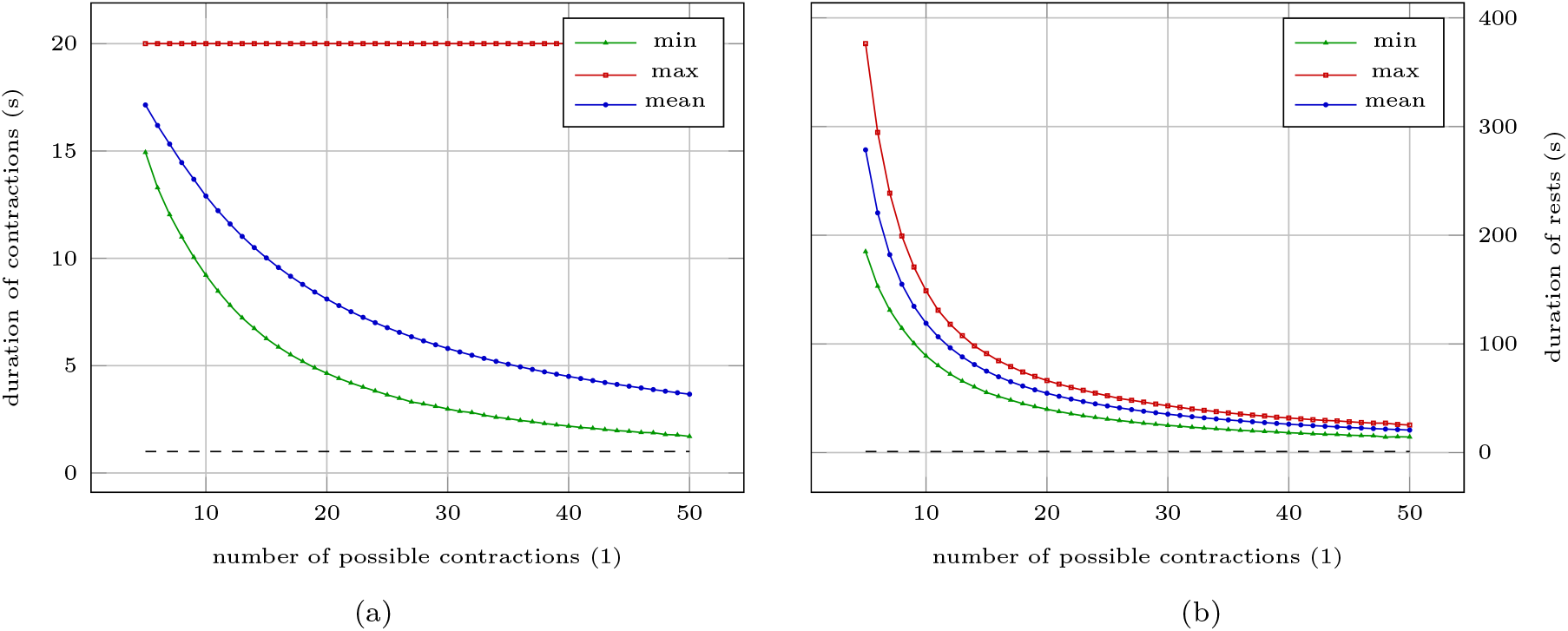
Dependency of the durations of contractions (a) and rests (b) on the number of possible contractions for Sessions D_5_ to D_50_. The horizontal dashed lines illustrate the 1 s mark. Increasing the number of possible contractions decreases the durations of contractions and rests of the computed solution.

Instead of choosing a minimum threshold intensity, we can emphasize higher loads by evaluating a weighting function on the integrand of the FTI. For demonstration purposes, we choose a quadratic weighting function for Session E. A similar approach has been used by Arandjelovic [6] to describe the hypertrophy stimulus of a resistance training set, although he used a sigmoid function, which can be interpreted as a smoothing of the constraint used in Session B. Figure 1g illustrates the model response obtained by simulating Session E.

For Session F, a similar quadratic weighting function can be applied to Session C. Figure 1h illustrates the model response obtained by simulating Session F.

### Fatigue-based goals

Effects of fatigue, e.g., metabolic stress or increased motor unit recruitment, have been attributed to trigger and/or positively influence muscle hypertrophy [39]. For Session G, we examine which loading scheme maximizes fatigue, defined as the accumulated loss of MVIC force over time. Figure 1i illustrates the model response obtained by simulating Session G.

For Session H, we maximize fatigue while ensuring a minimum threshold intensity of 80 % of baseline MVIC force. Figure 1j illustrates the model response obtained by simulating Session H.

In contrast to maximizing fatigue, it might also be desired to accumulate a certain amount of work while minimizing fatigue, e.g., during the tapering period before a competition. For Session I, we model such a scenario. Figure 1k illustrates the model response obtained by simulating Session I.

### TUT-based goals

Several authors have examined time-under-tension as a determinant of acute responses and longterm adaptations to RT (e.g., Burd et al. [12] or Schott et al. [40]). Therefore, for Session J, we maximize TUT while ensuring a minimum threshold intensity of 80 % of baseline MVIC force. Figure 1l illustrates the model response obtained by simulating Session J.

Session J does not take into account the duration of the contractions used to accumulate the total TUT. However, some author have reported different adaptations to short and long duration contractions with greater hypertrophy occurring after long duration contractions [40]. Thus, we weight the durations of contractions quadratically for Session K. All other settings are kept as in Session J. Figure 1m illustrates the model response obtained by simulating Session K.

### Durations of contractions and rests

Table 3 contains the minimum, the maximum, and the mean durations of the contractions and rests for all sessions plotted. To a certain extent, this allows to examine the real-life feasibility of the computed sessions.

**Table 3.** Minimum, maximum, and mean durations of contractions *δ_c_* and rests *δ_r_* for all sessions plotted. To a certain extent, this data allows to examine the real-life feasibility of the computed sessions.

| Session | $\min(\delta_c)$ | $\max(\delta_c)$ | $\text{mean}(\delta_c)$ | $\min(\delta_r)$ | $\max(\delta_r)$ | $\text{mean}(\delta_r)$ |
| --- | --- | --- | --- | --- | --- | --- |
| A | 19.21 | 465.46 | 60.54 | 1.96 | 8.76 | 6.49 |
| B<sub>70%</sub> | 6.24 | 33.28 | 11.41 | 28.63 | 45.64 | 38.11 |
| B<sub>90%</sub> | 1.62 | 9.13 | 3.04 | 33.02 | 56.96 | 46.83 |
| C | 3.71 | 6.06 | 4.11 | 28.90 | 51.81 | 45.72 |
| D<sub>5</sub> | 14.94 | 20.00 | 17.14 | 184.96 | 376.31 | 278.57 |
| D<sub>50</sub> | 1.70 | 20.00 | 3.67 | 14.38 | 25.36 | 20.75 |
| E | 16.10 | 62.36 | 26.06 | 7.15 | 25.63 | 22.86 |
| F | 3.08 | 6.54 | 3.52 | 39.36 | 73.22 | 59.45 |
| G | 1200.00 | 1200.00 | 1200.00 | 0.00 | 0.00 | 0.00 |
| H | 4.30 | 21.76 | 6.97 | 20.57 | 54.54 | 42.74 |
| I | 6.51 | 12.05 | 7.25 | 30.09 | 48.14 | 42.45 |
| J | 3.69 | 21.76 | 6.99 | 30.57 | 51.91 | 42.72 |
| K | 5.81 | 21.76 | 12.01 | 42.10 | 126.68 | 95.97 |

## Discussion

### Choice of training goals

In general, a model-based approach is limited by the predictive ability of the employed model and the available numerical solution methods. As mentioned, the model of Herold et al. [24] offers a phenomenological description of muscular fatigue for different loading schemes and does not directly link the RT input to a physiological adaptation of the trainee. Thus, when choosing the training goals, we are limited by key performance indicators accessible in the model. For this reason, we use assumptions from sport science about optimal training as objectives and constraints.

The three KPIs force-time integral, time-under-tension, and loss of MVIC force can readily be used in the optimal control problem formulations. Furthermore, we employ variants of these three KPIs to demonstrate how even slight modifications can change the structure of the solution. This highlights how important it is for exercise physiologists and sport scientists to identify the correct driving stimuli for adaptations in order to design optimized RT programs. Suitable physiological models would allow a more thorough search, e.g., by incorporating the build up of metabolites such as hydrogen ions and inorganic phosphate or by describing the activation of different fiber types.

### Structure of the computed RT sessions

While the resulting differences between the solutions might seem small at first, one should keep in mind that these differences accumulate during the course of an RT plan over weeks and months.

The results of Session D favor a higher number of contractions to accumulate more force-time integral in this scenario. This is in line with the solutions of most other sessions, in which all 25 possible contractions are realized. However, this is not the case for the solutions of Sessions A, F, G, and K. The results of Session A illustrate that the inclusion of rests is not beneficial during the beginning and the end of the session for this setting. To enable high contraction intensities, the solution of Session F consists of only 20 contractions. This is due to the fact that we weight the contraction intensities proportionally more than in the solution of Session C, where all 25 contractions are realized. The solution of Session G describes a sustained MVIC effort, which is caused by choosing the accumulated loss of MVIC force as training goal. The solution of Session K only realizes 12 contractions in order to enable longer contraction durations compared to the solution of Session J. This can be verified by comparing the mean contractions duration of Session J and K, i.e., 6.99 s and 12.01 s (see Table 3).

Except for the solutions of Sessions H, J, and K, all solutions consist exclusively of MVIC efforts. This was unexpected, as we anticipated that submaximal contractions might allow a greater accumulation of training volume due to them inducing less fatigue. It would be interesting to examine if such a behavior also occurs for dynamic constant external RT. The solution of Session H exhibits an interesting behavior as the inclusion of a minimum threshold intensity now favors submaximal contractions compared to the MVIC efforts of the solution of Session G. This is possibly caused by the longer contraction durations, which then contribute more to the accumulated fatigue. Session I exhibits the same behavior as the MVIC efforts reduce the time necessary to accumulate the desired FTI. The same holds for the solutions of Sessions J and K, where the submaximal contractions allow a greater time-under-tension. The submaximal contractions are all hold until failure. In case this is not desired, this could be included into the optimization problem as a constraint. If a minimum threshold intensity was chosen, the MVIC efforts are hold until this intensity is reached (see in particular Session B). Sessions C and F differ. Here, the contractions are terminated earlier as contractions with the minimum threshold intensity do not contribute to the chosen training goal. Session E demonstrates how a focus can be set on higher contraction durations without the use of a minimum threshold intensity.

As already noticed during the model development [24], the grouping of repetitions into sets is not supported by our results. Instead, the contractions are spread more evenly over the whole time horizon to allow a greater accumulation of training volume, i.e., force-time integral. This is a similar approach to variants of so-called cluster sets [45], which allow to increase training volume by breaking up the traditional set-repetition structure. Here, the algorithmic optimization of durations of contractions and rests provides a clear advantage over intuitive planning.

### Real-life feasibility of the computed RT sessions

To ensure the real-life feasibility of the computed RT sessions, several aspects have to be taken into account. First, the duration of the contractions may not be too short, as the trainees need time to develop MVIC force. Second, the duration of the submaximal contractions may not be too long, as the concept of task failure or limited work capacity is currently not implemented into the model [24]. Third, the rest periods between submaximal contractions may not be too short, as the model also does not account for a regeneration of work capacity.

Kawakami et al. [26] examined 100 intermittent MVIC efforts lasting 1 s followed by 1 s rest of the triceps surae muscles and reported no problems in executing this task. Table 3 and Figure 3 show that our solutions do not propose durations shorter than 1 s for contractions and rests. Although a different muscle group was used in the study of Kawakami et al. [26], their data demonstrates that such short intermittent contractions might be possible in general.

Yoon et al. [49] examined endurance times for sustained isometric contractions of the elbow flexors at 90 degrees joint angle and at 80 % of MVIC force. Although the experimental setup differed slightly compared to that of the experiments used for the model validation [24] (forearm horizontal versus forearm vertical to the ground), the mean endurance times of 25.0 s for men and 24.3 s for women are consistent with the maximum duration of 21.76 s of our solutions for Sessions H, J, and K (see Table 3). To the best of our knowledge, no prediction of endurance time or work capacity exists for MVIC efforts. Caffier et al. [16], for example, examined MVIC efforts of several muscle groups lasting 10 min and reported no task failure among the subjects. Thus, it remains to be validated experimentally if the solutions of Session A, E, and G, which contain sustained MVIC efforts of long durations, can be realized in practice.

Although several authors have examined the recovery of endurance times (see, for example, the work of Stull and Kearney [42] or Kroon and Naeije [27]) and work capacity (see, for example, the review by [25]), to the best of our knowledge, no model of their time course exists that fulfills the prerequisites postulated for use in an optimization context [24]. Furthermore, we are not aware of any experimental data that rejects the feasibility of the solutions of Sessions H, J, and K due to too short rests. If this should be the case, lower bounds on the durations of the rests could be incorporated into the optimal control problem.

### Limitations and future research

Our work is not free of limitations and several directions of future research are possible.

As no fully suitable mathematical model for the more commonly used dynamic constant external resistance (DCER) training is available, we are optimizing isometric RT sessions. Research shows that the transfer from isometric RT to dynamic performance is questionable [34]. Therefore, we discourage direct transfer of our findings to DCER or other forms of training. However, an extension of our approach to DCER training is straightforward once suitable models become available. The same holds true for extensions to other indicators of muscle fatigue (e.g., power, contraction velocity, or muscular endurance), multiple exercises, or longterm planning.

Moreover, we are using parameters obtained from the elbow flexors, as so far those are the only ones available. For this reason, a comparison between muscle groups or subjects is not possible at the moment. It would be intriguing to calibrate the model to different muscle groups and subjects and then examine how the resulting parameters affect the optimized RT sessions. Lievens et al. [30], for example, after analyzing fatigue and recovery patterns of MVIC torque of the knee extensors, conclude that individualizing training might be important to optimize performance. The authors used proton magnetic resonance spectroscopy to analyze muscle fiber typology of the gastrocnemius and then classify the subjects into a slow- and a fast-twitch group for which they expected different patterns. With a model-based approach, this classification could be formulated as a parameter estimation problem for which the necessary force measurements could be obtained in a single testing session [23]. Afterwards, RT sessions could be optimized individually as proposed in this work.

Last, we acknowledge that the model is validated with data from laboratory studies. Thus, we face the same problems as the original studies: the transfer from the laboratory to real-life RT needs to be verified experimentally.

## Conclusion

In this work, we demonstrate that a mathematical model-based approach could provide valuable impulses for practitioners and complement the predominant manual program design of loading schemes for RT. Although, the differences in the optimized sessions might seem small, one should keep in mind that those accumulate during the course of an RT plan over weeks and months.

As our approach is independent of the underlying model, we encourage researchers to develop and validate models, which are suitable for optimization and which connect the training input directly to training goals such as increasing strength and power, hypertrophy, or increasing local muscular endurance. This would extend the possibilities to set up the optimization problems and might furthermore help to identify the driving mechanisms for longterm adaptations. Then, we could exploit the full potential of our approach.

## Author contributions

JLH conceived the idea for this work. JLH conducted the literature research, performed the numerical experiments, and drafted the manuscript. JLH and AS discussed and edited the draft. JLH and AS approved the final version of the manuscript.

## Acknowledgments

JLH gratefully thanks Dr. Christian Kirches of the Institute for Mathematical Optimization, Technische Universität Carolo-Wilhelmina zu Braunschweig, Braunschweig, Germany for stimulating discussions on this topic.

